# *In vitro* effects on cellular shaping, contratility, cytoskeletal organization and mitochondrial activity in HL1 cells after different sounds stimulation. A qualitative pilot study and a theoretical physical model

**DOI:** 10.1101/2020.03.19.993618

**Authors:** Carlo Dal Lin, Claudia Maria Radu, Giuseppe Vitiello, Paola Romano, Albino Polcari, Sabino Iliceto, Paolo Simioni, Francesco Tona

## Abstract

Convincing evidence has documented that mechanical vibrations profoundly affect the behaviour of different cell types and even the functions of different organs. Pressure waves such as those of sound could affect cytoskeletal molecules with coherent changes in their spatial organization and are conveyed to cellular nucleus via mechanotransduction. HL1 cells were grown and exposed to different sounds. Subsequently, cells were stained for phalloidin, beta-actin, alpha-tubulin, alpha-actinin-1 and MitoTracker^®^ mitochondrial probe. The cells were analyzed with time-lapse and immunofluorescence/confocal microscopy. In this paper, we describe that different sound stimuli seem to influence the growth or death of HL1 cells, resulting in a different mitochondrial localization and expression of cytoskeletal proteins. Since the cellular behaviour seems to correlate with the meaning of the sound used, we speculate that it can be “understood” by the cells by virtue of the different sound waves geometric properties that we have photographed and filmed. A theoretical physical model is proposed to explain our preliminary results.

## 1. Introduction

There has been vast evidence that shows how cells are able to communicate with each other through light emissions^[1,2]^, electromagnetic signals^[3,4],[5]^ and mechanical / acoustic vibrations^[6,7],[8],[9]^.

In particular, it appears that mechanical vibrations profoundly affect the behavior of different cell types and even the functions of different organs^[10,11],[12],[13]^. Pressure waves such as those of sound could affect certain cells or their structures determining microvibrations, or even causing resonances, i.e. the synchronization of the biomolecular oscillatory patterns within cells^[14]^.

For example, it has been shown that bacterial cells are able to respond to specific single acoustic frequencies and are able to emit sounds^[15]^.

Moreover, it has been demonstrated that acoustic vibrations in the form of single frequencies^[16]^, noise or music, alter proliferation, viability^[17]^ and hormone binding ^[18]^ in human cells cultures and in animal models^[19,20],[21]^.

Ventura et al. have recently reported that human stem cells seem to respond to complex sound frequencies (melodic music, rhythm patterns and human voice) with different electromagnetic emissions measured with a Multi Spectral Imaging system^[22]^.

Furthermore, acoustic stimuli seem to be of paramount importance in guiding the spatial interaction between cells, thus influencing their individual and collective behavior^[23],[24]^.

It is very probable that all this evidence transmits, at a macroscopic level, the phenomena of cellular mechanotransduction, a discipline that focuses on how extracellular physical forces are converted into chemical signals at the cell surface. However, mechanical forces that are exerted on surface-adhesion receptors, such as integrins and cadherins, are also channeled very rapidly along cytoskeletal filaments and concentrated at distant sites in the cytoplasm and nucleus^[9,25,26]^, altering cellular genome activities^[27]^.

All the research conducted so far has mostly explored the possible biological effects of ultrasound or audible acoustic vibrations only in terms of cellular behavior, often using single or very simple frequencies; but what happens inside a living cell exposed to sounds? What could be the molecular effect of music, noise or of the words we hear or pronounce in our everyday life?

Since words and music are a compound of several sound frequencies, and are indeed mechanical vibrations, which can cause mechanical stress, it seems not odd to expect a direct molecular effect in our cells. The present work was thus designed to better understand the direct effects of acoustic vibrations in the form of music, noise and words in cardiac muscle cells *in vitro* culture in their contractility and their tendency to form cytoskeletal cellular networks in space.

Microtubules represent one component of the cytoskeleton and are composed of dimers of alpha e betta tubulin. Microtubules interacts with the other cytoskeletal proteins and it is responsible of the structure, shape of cell and in its movements. Microtubules interacts with other cytoplasmic organelles such as mitochondria and regulates their functions (metabolic energy)^[28]^. Microtubules system undergo in a rapid polymerization and depolymerization turnover by exchange their subunits (in the order of msec). In cardiac cells the interaction of microtubules system and mitochondrial play an important role in providing the bioenergy needs for the contraction of cardiac cells^[29]^. As microtubules, actin is involved in a dynamic process of polimerizzation-depolimerizzation. Thus, we focused our attention on these dynamic systems.

We exposed cultures of murine atrial cardiomyocytes (HL1) to 20-minute sound sequences. In particular, as an example of sounds with opposite characteristics, we used some meditative music, utterance, music with high frequencies, high rhythm and saturated sounds, a collection of urban traffic noises and human screams. A 20 minute repetition of single words: a sound that is repeated during meditation practices, and the sounds “Ti amo”, (I love you) and “Ti odio” (I hate you) were also used. These last sounds, as an example of words commonly used, with opposite meanings. We followed the cellular behaviour (contraction, vesicular trafficking, indirect cytoskeletal movements) directly in response to each different acoustic stimuli, acquiring a time lapse video with frames taken every 100 msec. Then, we analyzed any changes of the cytoskeleton by immunofluorescence technique. Finally, in order to explain the results, we also photographed and filmed the geometric characteristics of the waveforms of the sounds used.

## 2. Results

The HL1 cells were treated with different sounds for 20 min. The experiment was repeated for 6 times with the same outcome. The results were divided into two groups according to the stimulating or inhibiting effect of the sound (Supplements-Figure S2). Below we present the results for the vibration “Ti amo” and “Ti odio” as representative of the two categories (panels B and C of all the figures) compared to those of the control with spontaneous cell’s growth without any sound (panels A of all the figures). As depicted in the Figure S1, the other sounds we used produced similar effects. In all the figures, the sounds travel from right to left.

Control cells without any sound treatment portray casual organization of cells and normal expression of the different markers (panel A of all the figures).

HL1 cells were stained for alpha α-tubulin and MitoTracker^®^ (MTr). Following the “Ti amo” sound exposition, the expression of α-tubulin increased drastically and the mitochondrial marker decreased considerably (Fig. 1-B) while after the “Ti odio” sound the cells decreased sharply the expression of the same markers (Fig.1-C). Furthemore, tubulin distribution in Fig. 1-B seems to be organized as a net of filaments perpendicular and parallel to sound waves. In Fig. 1-B we can notice how tubulin forms “bridges” between cardiomyocytes, an important aspect of cell to cell communication and contractility^[49]^. After the “Ti odio” sound exposition (Fig.1-C), cells show spatial disorganization and their number decreased probably due to its “explosion” (Supplements-movie S1 series).

**Figure 1.**
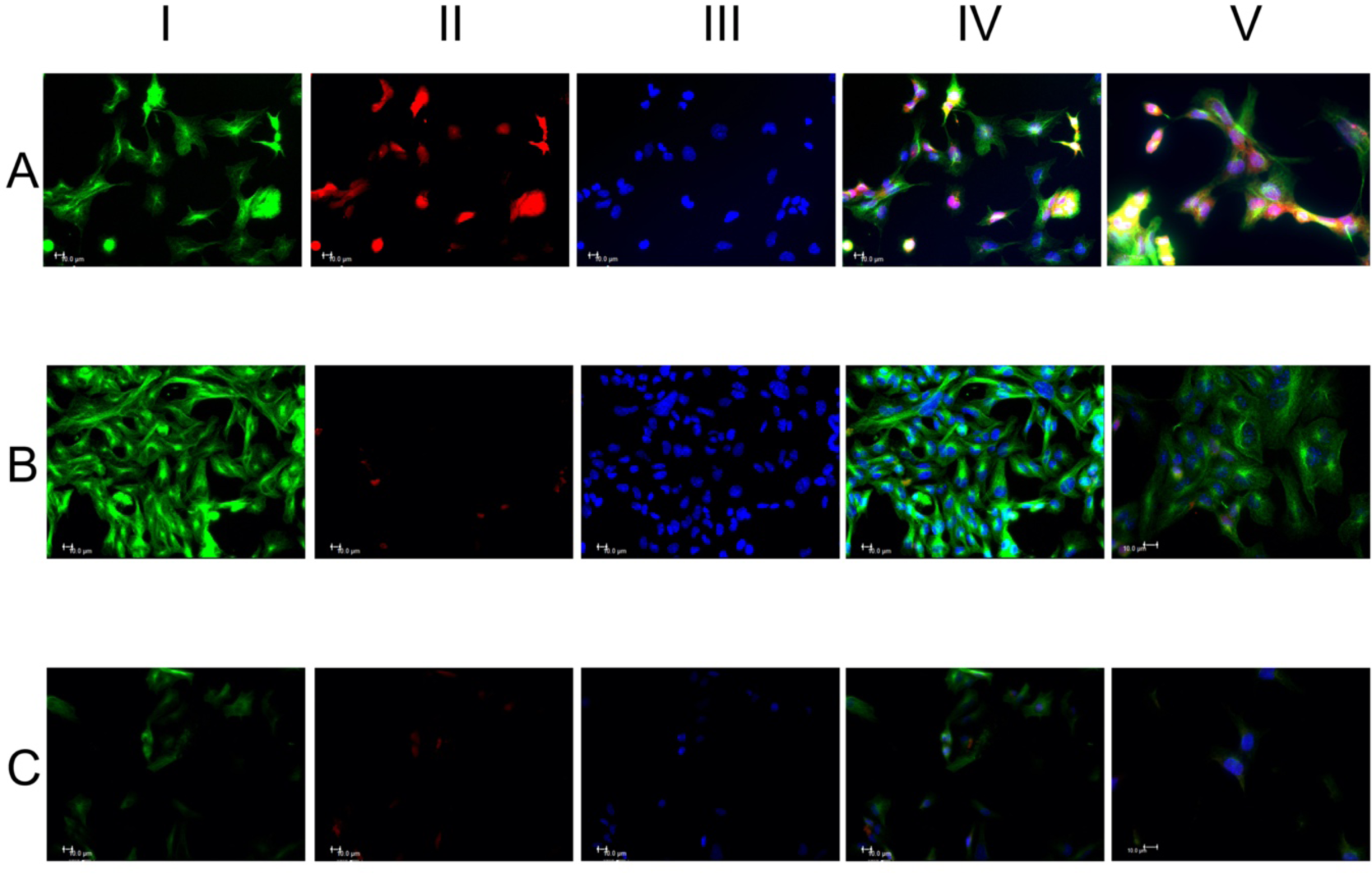
Alpha tubulin (α-tubulin) and MitoTracker (MTr) stained after treatment with different sounds for 20 min (representative images). The cells received the sound from right to left. HL1 cells were stained with α-tubulin (green fluorescence) column I, MTr (red fluorescence) column II and the nuclei were stained with Hoechst 33258 (blue fluorescence) column III. Merge of the three fluorescence, column IV. Representative merge fluorescence images at 100x/1.4 oil objective, column V. Panel A, control cells without any sound treatment showing casual organization of cells and normal expression of the different markers. Panel B, the expression of α-tubulin increased and the mitochondrial marker decreased. Cells with more orderly spatial organization with perpendicular tubulin distribution and parallel to sound waves. The nuclei appear with more regular size. Analysis with a greater objective (column V, panel B) highlights the filamentous and granular cytoplasmic expression of tubulin. In the image tubulin forms “bridges” between two cardiomyocytes which is important in the cell to cell communication (asterisk). Panel C, the HL1 cells show a spatial disorganization and the number of cells decreased. Moreover, the different markers expressions markedly decreased. The low number could be due to the “explosion” of cells with cytoskeletal fragmentations (asterisks). Scale bar 10µm.

In Fig 2. HL1 cells were stained with phalloidin (F-actin) and β-actin. The expression of both F-actin and β-actin increased after the “Ti amo” sound, (Fig. 2-B) while the cells decreased the expression of the same markers after the “Ti odio” sound (Fig. 2-C), as confirmed in Table I that describes their mean intensity fluorescence in three different ROI. The augmented expression of these markers in Fig. 2-B may indirectly reveal an increased cell activity and motility^27^ as shown in movie S2 series-supplements.

**Figure 2.**
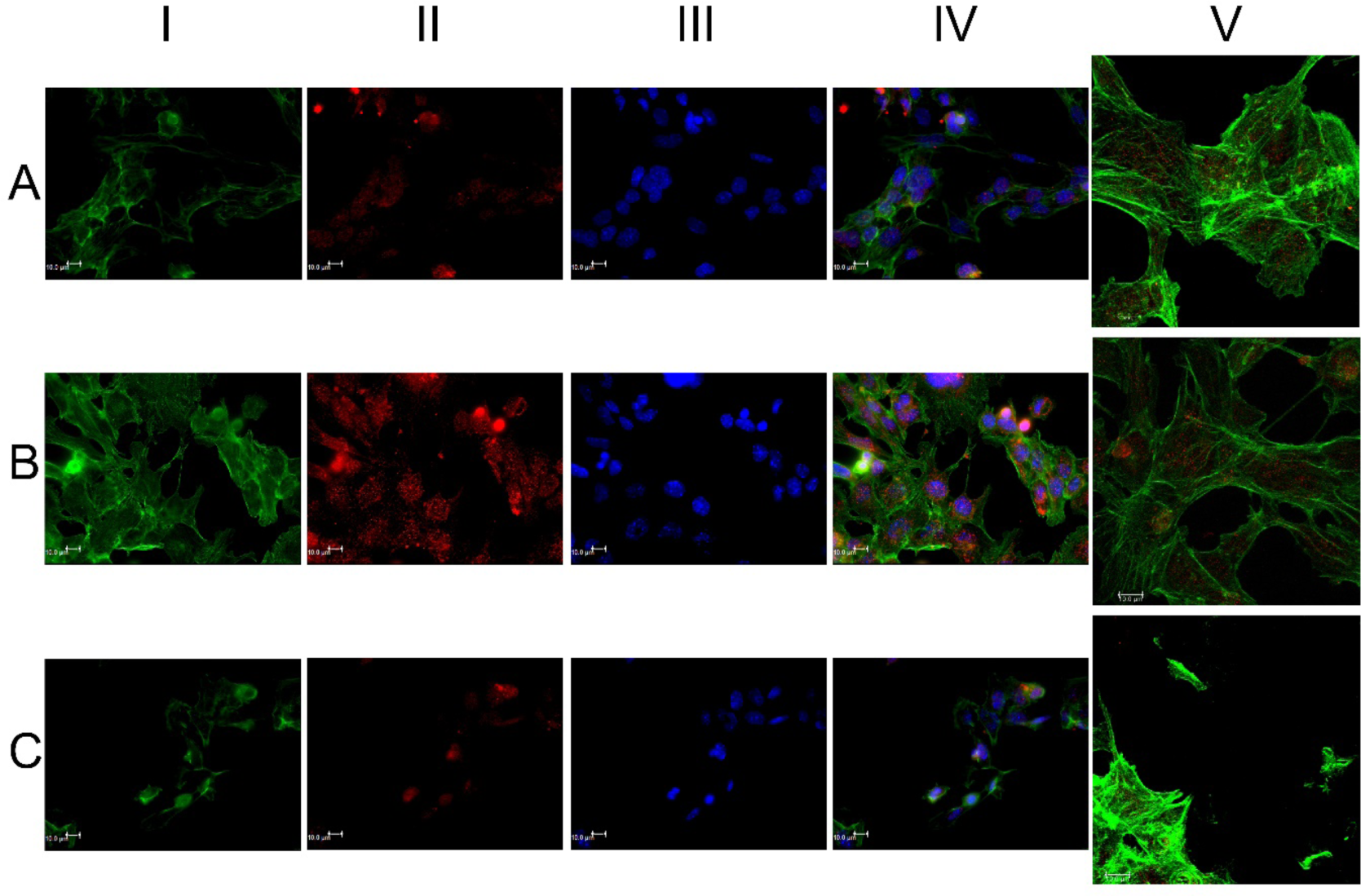
HL1 cells were stained with phalloidin 488 conjugated (green fluorescence) column I, β-actin (red fluorescence) column II and DNA with Hoechst 33258 (blue fluorescence) column III. Overlay of the three fluorescence images, column IV. Representative merged images of F-actin and β-actin with confocal microscope (100x/1.4 oil objective), column V. Panel A, control cells without any sound stimulus and normal expression of the different markers. Panel B, increased of both expressions of the two markers. Peripheral and filamentous expression of phalloidin and with a spatial organization as descripted in Fig. 1. Panel C, decreased number of cells with markedly fragmentation of cells (asterisks). Evident reduction of the expression of the different markers. Scale bar 10µm.

In Fig 3. cells were stained with F-actin and alpha-actinin-1. In controls (Fig.3-A), alpha-actinin-1 localization seems to be more perinuclear. After the “positive” sound exposition, its distribution is more diffused (Fig.3-B) while after the “negative” sound its expression is strongly decreased. We want to highlight the alpha-actinin-1 builds bridges between F-actin filaments^27^, which is another aspect that could further explain the increase in cell contractility shown in movie S2 series-Supplements.

**Figure 3.**
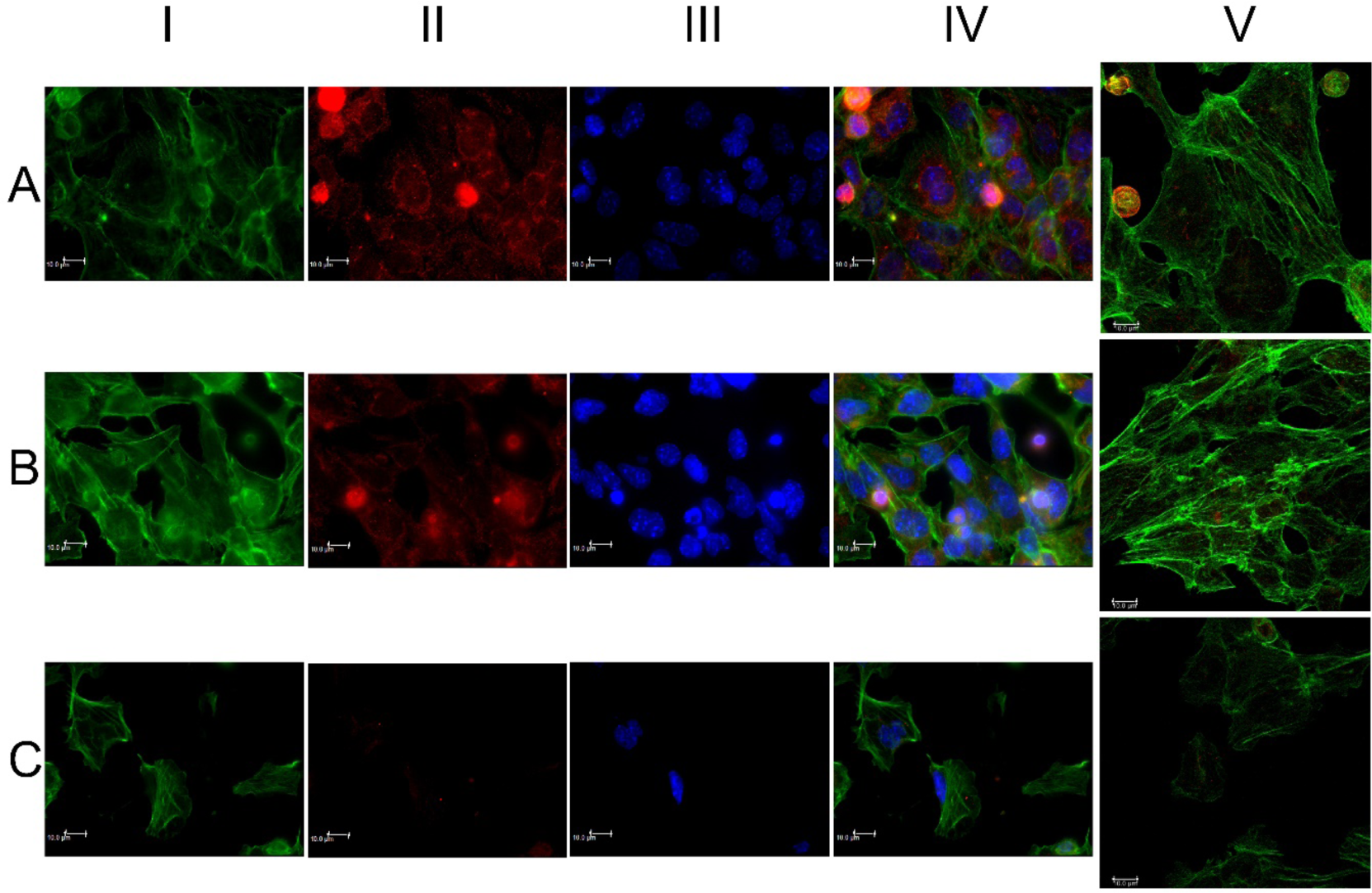
Cells stained with phalloidin and alpha-actinin-1(green and red fluorescence, column I and II, respectively) and DNA stained with Hoechst 33258 (blue fluorescence). Panel A, perinuclear alpha-actinin distributions. Panel B, major cellular organization and diffuse alpha-actinin distribution. Panel C, evident cellular fragmentation with loss of the normal morphology of the cell with nucleus towards the peripheral part (white asterisks). Decrease or absence of markers expression with following loss of cytoskeletal organizations (asterisk orange). Scale bar 10µm.

Table I quantitatively describes the molecular expression changes according to the sounds used.

**Table I.** Mean fluorescence intensity of 3 different Regions Of Interest (ROI) taken in 3 different images of the same marker.

|  | A | B | C |
| --- | --- | --- | --- |
| **Phalloidin** | 17.8±4.3 #* | 30.3±5.4 #● | 6.1±2.9 *● |
| **Beta-actin** | 1.5±0.3 #* | 7.4±1.9 #● | 2.7±0.6 *● |
| **Alpha-actinin-1** | 1.5±0.2 #* | 3.1±0.7 #● | 1.0±0.2 *● |

#Mann-Whitney for independent samples A vs B $p<0.05$
\*Mann-Whitney for independent samples A vs C $p<0.05$

## 3. Discussion

Qualification is not quantification. We are describing a proof of concept study and we need to make an important clarification to understand our results: our aim was to document any *qualitative* difference in cell behavior and cytoskeletal characteristics without focusing on *quantitative* assessments that will be the subject of further studies. Every biological phenomenon has qualitative and quantitative aspects of equal and complementary importance. Indeed, a quantitative expression variation of a subcellular component does not inform us about any possible conformational change or of a different spatial localization that could, even with the same quantity, justify a modification of cellular behavior. Any quantitative difference does not constitute a condition neither sufficient nor necessary to explain a biophysical phenomenon or, in our case, the interaction between sound vibrations and *in vitro* cells. We consider more appropriate to integrate quantitative studies with qualitative observations, from which we decided to start replicating them for 6 times.

Our results seem to highlight a different cell growth pattern depending on specific sound stimuli, with a different expression/organizzation of cytoskeleton proteins and mitochondrial activity^[28]^. The cellular behaviour of HL-1 cells also appears to change as a function of the sound stimulus that impacts on their surface: some biomechanical acoustic stimulations enhance contractility and increase the vesicular traffic within them, while others hinder it, reaching cell lysis. We speculate that this could be related to a transcriptional stimulation effect, based on the observation of Wang et al^[27]^.

The present observations and the videos in the supplementary material show how HL-1 cells respond in a highly sensitive fashion to vocal sounds and music or noise with opposite and, in hindsight, “predictable” results based on the type or the meaning of the acoustic stimulus itself.

The sounds used in this work are obviously composed of a spectrum of peculiar acoustic frequencies that propagate in a liquid medium (such as the culture medium used) by drawing different geometric shapes (movie S3-supplements) as already demonstrated by Chang et al^[30]^.

We assume that the meaning of the sounds we used could be encoded in their specific geometric form, and that the wave trains characteristic of each acoustic vibration can: 1) directly modulate the cytoskeleton morphology^[26,31]^ by penetrating signals of mechanical stress or relaxation up to the nucleus, thus interfering favourably or not with the trafficking of organelles (such as mitochondria^[32]^), of cellular micro-vesicles^[33]^ and influencing protein oscillatory motions normally present in the cytoplasm^[34]^; 2) arrange the traffic and the electromagnetic interaction^[35],[36]^ between the molecules dissolved in the cytoplasm, regulating processes crucial to cell survival^[37,38]^ according to the mechanisms exemplified in the supplementary animations (movie S4-supplements). It is not surprising that there may be an intimate relationship between trains of acoustic waves and proteins^[37]^, given that the proteins in the human body vibrate in different patterns like the strings on a violin or the pipes of an organ^[39]^ and may interact with each other even through quantum interference^[40]^.

On one hand, it is known that acoustic stimulations deform^[41,42]^ and are conveyed^[26,43–46]^ by the cytoskeleton to the nucleus^[27]^ and on the other, the tubulin monomers exhibit different oscillatory and assembly modes^[47]^ depending on the frequencies that go through them^[3]^. Furthermore, microtubules seem to be able to capture *phonons*, i.e. “quanta of sounds” or vibrational / mechanical energy packages, within them, energy that would then be used in different cellular processes, including cell division and movement^[48]^. In addition, it seems that *phonons* play an important role in regulating the structure and stability of a crystalline system^[49]^: further studies will be needed to extend this consideration to a complex system like a living cell, whose membranes, DNA, proteins and cytoskeleton seem to present analogies with a crystal lattice.

We remember that this is only a qualitative pilot study and we specify that the sounds and words used in this work are only an example of possible vibrations that we emit or to which we are exposed every day and of which we wanted to see a possible cellular biomechanical effects. Then, further studies will be needed to evaluate the complex phonological features of different sounds, music and words, pronounced by different people in different languages and cultures. Moreover, our results refer to “Ti amo” and “Ti odio” just as vibrations. Further studies will be needed to establish if the meaning that we assign to a specific sound is only a convention or related to its physical/geometrical mechanical properties.

In Appendix-1 we hypothesize a theoretical physical model at the basis of our results which could be very important for future research.

It is really intriguing to note that, as well as a scream or sound conveying in a geometric waveforms vibration (“Ti odio”) seem to induce cellular stress, as revealed by James Gimzewski with an atomic force microscope^[50]^ that produces a sound of a scream.

## 4. Materials and Methods

In Supplements-Figure S*1* it’s possible to see the laboratory set up for the experiment.

### Cell culture

The murine atrial cardiomyocyte cell line HL1 was obtained from the laboratory of William C. Claycomb (New Orleans, USA). The cells were grown in Claycomb medium supplemented with 10% fetal bovine serum (FBS, cat # F2442), 0.1 mM norepinephrine, 100 U/ml penicillin, 100 µg/ml streptomycin and 2 mM L-glutamine (all reagents purchased from Sigma-Aldrich, Italy) at 37°C in a humidified 5% CO2 incubator. The cells were seeded (2×10^5^ cells /ml) in 24-well culture plates containing a glass coverslip in each well coated with 25 µg/ml fibronectin/0.02 % w/v gelatine solution (Sigma-Aldrich). The culture medium was changed every 3 days with fresh medium.

At 70-80% of confluence HL1 cells were exposed to 20-minutes of sound sequences. We used this specific time period because we wanted to begin to study *in vitro* any possible direct effect of sound that could help to explain the results that we are obtaining *in vivo* in other researches^[51]^.

We choose atrial cardiomyocytes for their contractility, in order to try directly to visualize with the Time-lapse technique (movies S-1 and S-2) possible changes of contraction in response to the mechanical stress related to the sound waves used. After seeing a differential behavior with specific sounds, we proceeded with the qualitative molecular analyzes, repeating the experiment 6 times, always with the same results.

### Time-lapse images

Cells at 80% confluence were analysed in live imaging using a Leica DMI6000CS microscope (Leica Microsystems, Wetzlar, Germany) equipped with an incubator, with a temperature-controller set at 37 °C and in a humified 5% CO2 atmosphere. The cells were acquired every 100 or 160 msec for 20 min using a phase contrast (Ph) or a differential interference contrast (DIC) at 20x/0.4 or 40x/0.6 objectives respectively. Images were acquired using a DFC365FX camera and using the Leica Application Suite (LAS-AF) 3.1.1. software (Leica Microsystems). Some time-lapse experiments were performed in the presence of 250 nM MitoTracker^®^ Red (Thermo Fisher Scientific, USA) probe for 20 min, a specific mitochondrial marker.

### Immunofluorescence and confocal microscopy

Immediately after the treatment of the cells with the different sounds, the cells were washed twice with phosphate buffered saline (PBS, 137 mmol/l NaCl, 2.6 mmol/l KCl, 8 mmol/l Na_2_HPO4, 1.4 mmol/l KH_2_PO_4_, pH 7.4) and fixed in 2% paraformaldehyde for 20 min, permeabilized with 0.5% Triton X-100 in PBS for 15 min and treated with 0.05 mol/l NH_4_Cl for 15 min. All steps were performed at room temperature. Subsequently, cells were stained with the following antibodies; mouse anti-alpha-tubulin (α-tubulin) (EXBIO Praha, Czech Republic) diluted 1:500, Alexa Fluor 488-Phalloidin diluted 1:150 to visualize F-actin and rabbit anti-alpha-actinin-1 (Thermo Fisher Scientific, USA) diluted 1:25, all the antibodies were incubated for 1h at 37°C. Rabbit anti-beta actin (β-actin) (Thermo Fisher Scientific, USA) diluted 1:200 was incubated over night at 4°C. The cells, after incubation with the primary antibodies, were stained with the following specific secondary antibodies; 1:100-diluted fluorescein isothiocyanate (FITC)-conjugated goat anti-mouse IgG (Chemicon International, Billerica, MA,USA), 1:200-diluted Alexa Fluor 594-labelled goat anti-rabbit IgG (Life Technologies). The primary and secondary antibodies were diluted in PBS containing 0.5% bovine serum albumin (BSA). Secondary antibodies were also used in the absence of primary antibodies in order to assess non-specific binding. For the immunofluorescence analysis, the cell nuclei were labelled with 1.5 µg/ml Hoechst 33258 (Sigma-Aldrich) for 20 min at room temperature. Finally, the slides were mounted with Mowiol anti-fade solution (Sigma-Aldrich).

Leica DMI6000CS fluorescence microscope (Leica Microsystems) was used and samples were analysed with differential interference contrast (DIC) and fluorescence objectives. Images were acquired at 40x/0.60 dry objective magnification. Images were acquired using a DFC365FX camera and using the Leica Application Suite (LAS-AF) 3.1.1. software.

The same samples were analysed by a confocal microscope TCS SP8 (Leica Microsystem) with a z-interval of 1.5 µm using a 100x/1.4 oil immersion objective (image size 1024×1024 pixel). Images were acquired with the above described camera and software.

Finally we measured the mean gray scale values (mean fluorescence intensity) of the markers phalloidin, beta-actin and alpha-actinin-1 using three different Regions Of Interest (ROI) taken in three different confocal images of the same marker, using the specific statistic function of the LAS-AF 3.1.1. software (Leica Microsystems). Mean fluorescence intensity data were compared using Mann-Whitney test for independent samples.

## Supplementary Materials

**Figure S1.**
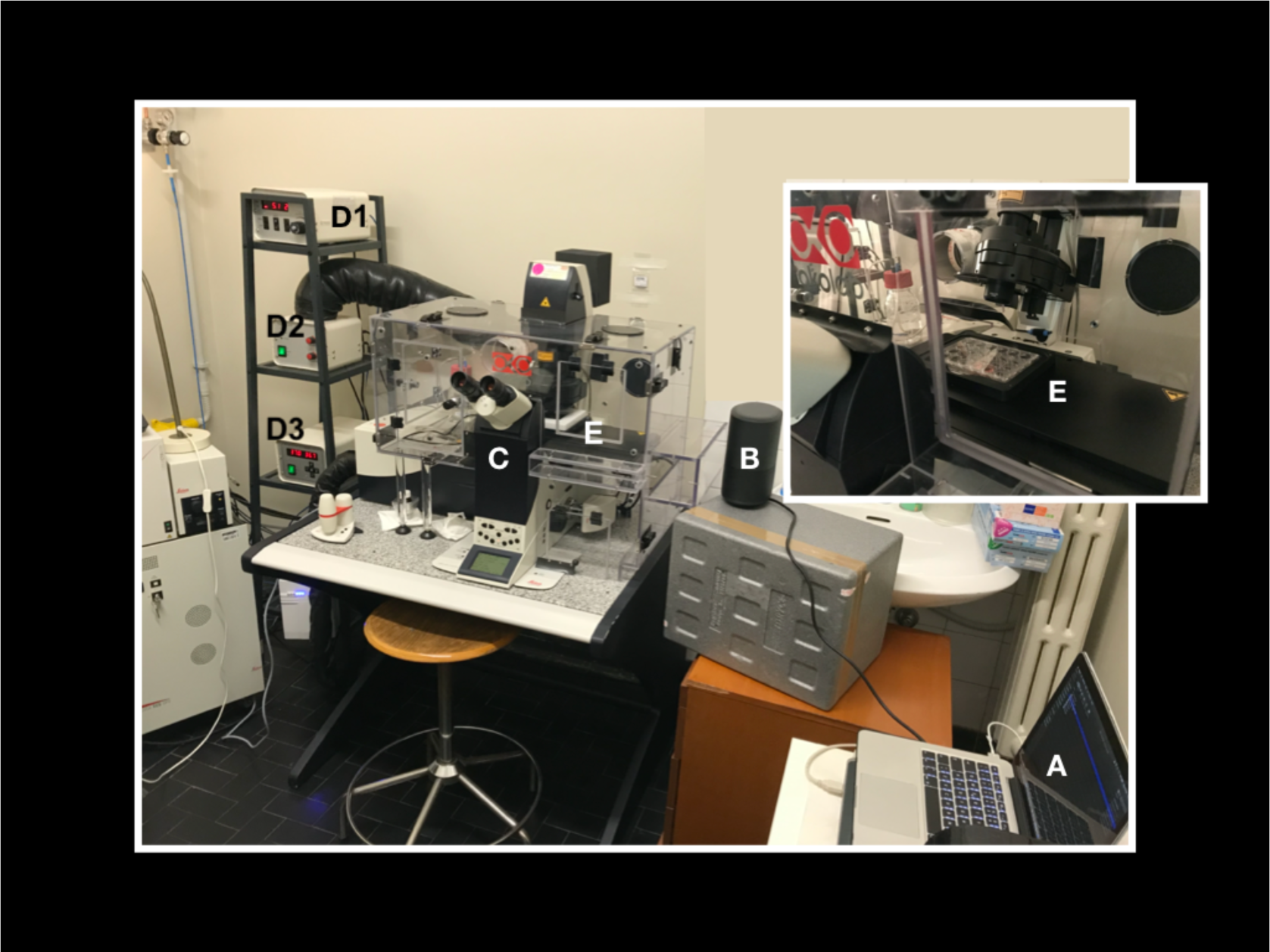
Our laboratory set up for the experiment. A laptop computer with the audio tracks (A) is connected to an amplifier (B) positioned nearby the microscope (C) and its incubator (E). With a phonimeter we calibrated the volume of the amplifier so that the intensity of the sound in the point where the cells are positioned (E) is about 60 dB. which the intensity corresponds to a normal vocal conversation. (D1) CO2 controller, (D2) heating unit, temperature control unit (D3). We want to specify that also “control cells” were exposed to the speaker plugged to energy without any sound produced in order to exclude that our findings could be the result of any small vibration produced by the speaker itself instead of the sound. Finally, the sounds “Ti amo”, “Ti odio” and the sound used in meditation were pronounced by 3 different people, giving the same results.

**Figure S2.**
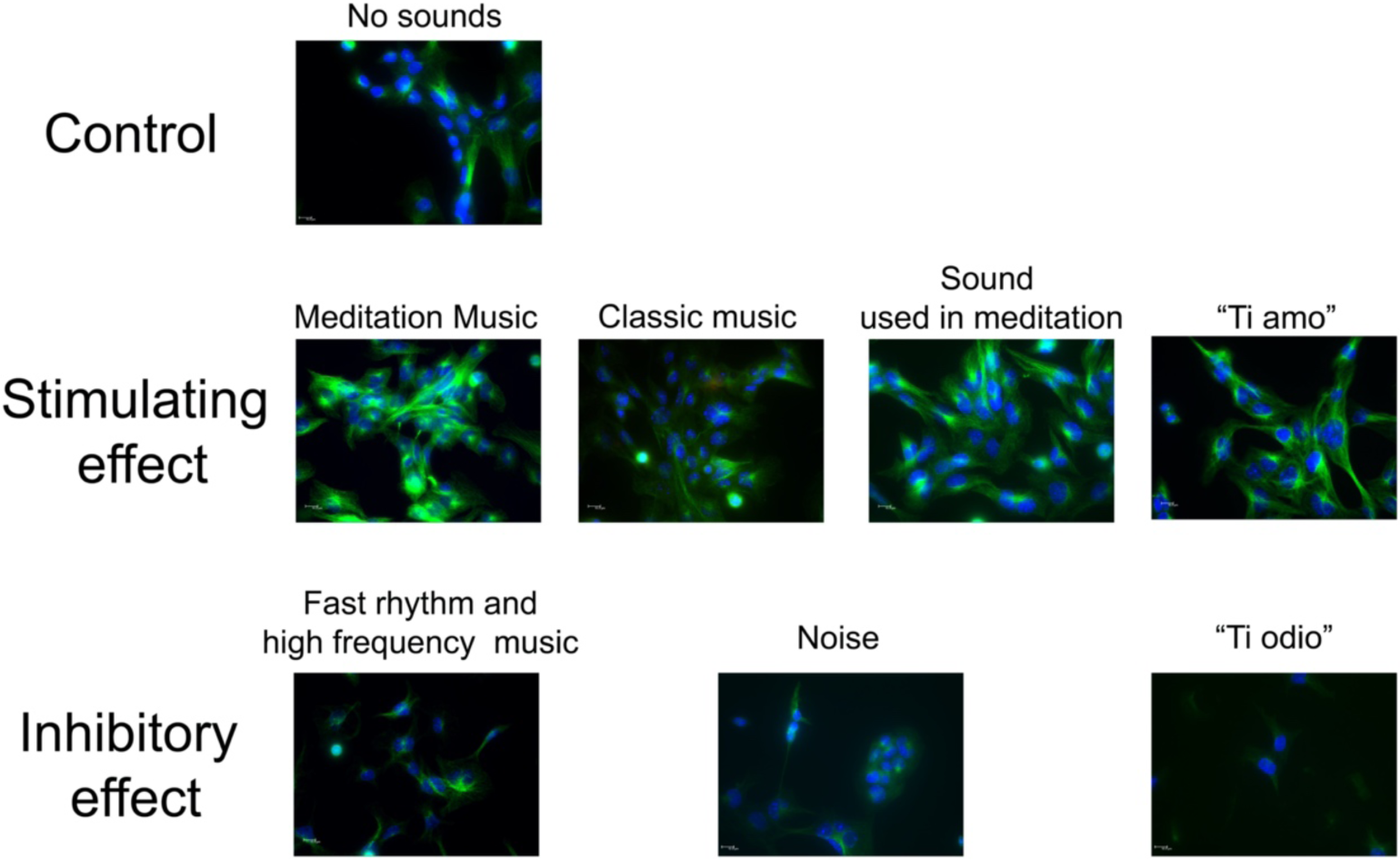
Alpha tubulin (α-tubulin) stained after treatment with different sounds for 20 min. The cells received the sound from right to left. HL1 cells were stained with α-tubulin (green fluorescence) and the nuclei were stained with Hoechst 33258 (blue fluorescence). The images show the merge of the two fluorescence at 63x/1.4 oil objective. In the first line, control cells without any sound treatment showing casual organization of cells and normal expression of the different markers. The middle line shows the stimulating/organizing effects of meditative and classic music, of a sound used in meditation and “Ti amo” sound. It is possible to observe the increased expression of α-tubulin. The third line depicts the inhibiting effects of high speed and high frequency music, noise (a collection of traffic noises and screams) and “Ti odio” sound. It is possible to observe the decreased expression of α-tubulin and the reduced number of cells. Scale bar 10µm.

**Movie S1**. Time-lapse video (images taken for 20 minutes every 160 msec) of HL-1 cardiomyocytes. A) controls, no sounds. B) “Ti amo” sound exposition. Please notice the increase in cell’s motility and in the inner vesicular trafficking, compared with the control. C) Please notice cellular explosion during the “Ti odio” sound exposition. In some frames, the images may seem a bit fuzzy: it is a normal consequence of shooting in 2D, a 3D structure while moving.

**Movie S2**. Time-lapse video (images taken for 20 minutes every 100 msec) acquired on the same field. This video shows the behavior of the same HL-1 cardiomyocytes during 20 minutes of spontaneous growth/activity without any sounds (A) followed by the “Ti amo” (B) and the “Ti odio” (C) sound exposition. Please notice the increase in cell’s contractility and the increased number of cells that start to beat in B. Finally, observe the decreased and asynchronous cell’s contractility in C and the cessation of the beating of some cells. In some frames, the images may seem a bit fuzzy: it is a normal consequence of shooting in 2D, a 3D structure while moving.

**Movie S3**. The video shows the different geometric shapes of the wave trains linked to the words “Ti amo” and “Ti odio” while traveling in water. A drop of water was placed in a Petri dish placed above a vibro-acoustic plate (VIBBRO srl-Italy) connected to a computer. The audio files are emitted by the PC and the plate vibrates according to the specific frequencies of the sound, describing different geometric patterns both on the surface and on the thickness of the aqueous medium. This could mimic what occurs inside a cell and how all the molecules inside it oscillate. We did not use this platform during the experiment (please see fig.S1). This platform served only to show how a vibration can spread in a watery environment, similar to the cytoplasm.

**Movie S4**. Acoustics and vibration animations (courtesy of Dr. Daniel Russel, Pennsylvania State University): the video shows how a particle placed in an aqueous environment (such as the cytoplasm) can move and oscillate (in compression and rarefaction cycles) as a function of the longitudinal and transversal component of the mechanical wave (sound) that passes it. We speculate that this movement could influence and regulate the interactions between the intracytoplasmic molecules and cell to cell communication. The different frequencies that make up a sound can travel compact in the case of a non-dissipative medium or separate into wave trains with different speeds in the case of a dissipative medium (such as the cytoplasm). The proteins polymers can undergo specific torsional deformations that could be influenced by mechanical waves such as sound waves.

## Author Contributions

conceptualization, CDL; methodology, CDL and CMR; formal analysis, CDL and CMR.; investigation, CDL and CMR.; resources, SI, PS and FT; data elaboration, CDL and MCR.; writing—original draft preparation, CDL and CMR; writing—review and editing, SI, PS and FT; supervision, SI,PS and FT; project administration, SI,PS and FT; funding acquisition, SI, PS and FT. Appendix-1: conceptualizzation GV and CDL; formal anlysis of sounds and graphs preparation GV, PR and AP, writing GV.

## Ethics approval

(not applicable).

## Availability of data and material

(not applicable)

## Code availability

(not applicable)

## Acknowledgments

we would like to thank prof. Daniel A. Russel for the acoustics and vibration animations contained in supplements series-video 4. We acknowledge Vibbro srl for having created and made available the vibrating platform with which we have visualized the sounds propagating in water; we thank the Entolé staff for the recording of audio files; the meditation music used in this study was composed by Paolo Spoladore.

## Appendix 1

*In vitro* effects on cellular shaping, contratility, cytoskeletal organization and mitochondrial activity in HL1 cells after different sounds stimulation. A qualitative pilot study and a theoretical physical model.

### The theoretical physical model

In this Appendix we discuss the effects on cells and their components that are induced by different sounds, pressure waves and mechanical vibrations, which we will refer to generically as ‘sounds’. We will resort to results reported in Refs.^[1,2,3]^. We call ‘positive’ sounds those producing ordering, formation of protein ‘bridges’, cell-to-cell correlations, strengthening of the links in the cytoskeleton network, etc., resulting in an augmented biological activity. The ‘negative’ sounds induce instead negative response, i.e. the inhibition of the ordering among biological components, sometimes presenting even the cell destruction (cell ‘explosion’)) (see Figs. 1-3, S2, Movies S1 and S2. See Figs. A1, A2, A3 for the representations of three sound signals).

A characteristic feature of quantum field theory (QFT) is the distinction between the symmetry properties of the states of the system and those of the dynamical equations. When the symmetry properties of the system ground state are not the same ones of the dynamical equations, we have the phenomenon of spontaneous breakdown of symmetry (SBS) which manifests itself in the generation of ordering (‘spontaneous’ means that the system dynamics selects self-consistently the system ordering). The formation of ordered patterns is due to long range correlations among the system constituents. The quanta associated to such long range waves, called the Nambu-Goldstone (NG) boson quanta, condense in the system ground state ^[4,5,6]^.

In biological matter, macromolecules are characterized by the specific electric dipole moment of their constituent units. Macromolecules and cells are embedded in the water bath. Water molecules, which constitute the large majority in number of the present molecules are also characterized by their specific electric dipole moment. Since dipoles may be oriented in any direction, the basic symmetry is the rotational spherical symmetry. The action of some external input, e.g. sounds, may trigger the breakdown of the dipole rotational symmetry by producing dipole ordering, namely their alignment along a preferred direction, or their in phase oscillations. The state of the system will be then characterized by the non-vanishing polarization density *P(x,t)* which provides a measure of the ordering (the electret) and it is therefore called “order parameter”. The NG quanta of the long range correlations responsible of the ordering are called dipole wave quanta (dwq) ^[1,2]^. The generation of the dipole long range correlations facilitates the occurrence of short range interactions, such as Van der Waals interactions, H-bonding, etc. by reducing the randomness of the molecular kinematics so to enhance the observed high efficiency of metabolic reactions. Long range correlations allow the collective, *coherent* behavior of the system components which manifests itself in the macroscopic scale behavior of the observed cellular structures and cell network.

Beside the sound induced SBS in the cell environment and in the water bath where the cells are embedded, there is another remarkable effect produced by sounds on quasi-unidimensional structures, such as protein chains. The energy supplied by the sounds hitting the extremity of a protein molecular chain may produce indeed a deformation in the chain turning into oscillations of the electric dipoles of the chain units. The conformational change, coupled to the chain molecular dipoles, traveling over the chain as a localized deformation is ruled by the nonlinear Schrödinger equation. It is known as the Davydov soliton, or Davydov solitary wave, so called since it remains stable under collision with other similar waves ^[7,8,9]^.

The energy necessary to excite the Davydov soliton is of the order of *0.25* eV, which is the energy released by ATP hydrolysis at the origin of a protein chain ^[1,2,3,7]^. In addition to the possibility that the sound directly couples to the chain, we may also suppose that the energy supplied by the sound triggers the ATP reaction and this one in turn triggers the soliton formation on the protein. It is not relevant to our analysis which one of these possibilities is actually realized. In both possible cases, the response of the protein chain to the sound is the propagation in the form of the Davydov solitary wave of the *coherent* localized deformation, i.e. of *in phase* dipole vibrational modes. In many-body physics the ‘coherent dynamics’ is indeed the one ruling the “in phase motion” of a collection of elementary components ^[1,2,3,4,5,6]^.

We list few of the properties of the dynamical scenario implied by the soliton:

i. The solitary wave has a ‘nondissipative’ propagation. This means that contrarily to the usual (non-solitary) wave motion, the soliton does not lose (or loses very little amount of) its energy during its traveling on the protein chain and can propagate over large distances. Since the discreteness of the chain may produce periodic potentials, the soliton velocity may be a function of time ^[8]^. The energy supplied by the sound to the system may remain thus ‘trapped’ in the solitary wave and it is then transferred along the protein chain (almost) without dissipation. In the absence of such a dynamical mechanism an excess of sound energy supplied to the cell, with a frequency spectrum unable to resonate with the molecular oscillatory motion of the cell constituents, might lead to the excessive warming up of the cell, possibly leading to its explosion, as observed in the case of ‘negative’ sounds.
ii. The oscillations of the chain molecular dipoles propagating with the soliton produce an electromagnetic (e.m.) radiation. Such a radiation constitute an endogenous source of e.m. field and will then break the rotational symmetry of the surrounding water molecular dipoles with their consequent coherent oscillation and orientation. Spectroscopic observations under ambient conditions have indeed shown ^[10]^ that hydration water forms a superstructure of the macromolecule (DNA, in the experiment of Ref. ^[10]^) with a strict interdependence between deformations of the macromolecule chain and its water environment.
iii. A soliton is able to trap an electric charge in its motion (electrosoliton) ^[7,8,9]^. This may then result in a nondissipative electric current flowing (with the soliton) over the protein chain. The radiation emitted according to the Maxwell equations, together with the one emitted by the chain dipole oscillations (point ii)), synchronizes with the radiation emitted by solitons on the other proteins; they might then travel with the same velocity and emit radiation at the same frequency ^[9]^.
iv. The soliton traveling over an infinitely long chain is stable (infinite life-time). Protein chains have, however, finite length. The Davydov soliton has then a finite life-time. It decays once in its propagation has reached the end of the chain and will then release its trapped energy in the surrounding water bath. Such an energy will not produce an increase of the bath temperature since the energy will propagate in it not in a diffusive way, but in wave form because the bath has been brought in the electret (ordered) state by the same soliton (point ii)) before its decay.

The supply of energy to the cell by the sound can be regarded as a “feeding” process of the cell. As described above, such a feeding process triggers the spontaneous breakdown of the rotational symmetry of surrounding water dipoles with the generation of the water electret state. The short life-time of the water electret state suggests to us the cyclic process picture, by which, in order that the functional biological activity related to the ordered cell state may continue, the cell needs to be fed again by an external resonating agent when the electret state vanishes, and so on, in a cycle, indeed.

Summing up, the emerging picture seems to fit with the experimental observations reported in text for the cases of ‘positive’ and ‘negative’ sounds. The ‘negative’ effects on the biological matter may be attributed to the lack of the dynamic processes described above (cf. Figs. 1-3, S2, Movies S1 and S2).

The sound induced ordering processes considered above generate in turn another remarkable effect. It is known that the e.m. field propagates in a self-focusing fashion through an ordered medium and acquires a finite mass in its propagation (the Anderson-Higgs-Kibble (AHK) mechanism) ^[5,6]^. This provides the understanding of many phenomena in condensed matter physics and elementary particle physics. In our case, the mass *M* acquired by the e.m. field turns out to be a function of the water polarization density *P(x,t), M = M(P), M(P)= 0* at *P=0*, and the propagation of the field occurs in a filamentary-like fashion, as through channels piercing the ordered medium (self-focusing) ^[11,12,13]^. The Maxwell propagation in spherical waves is restored at *P(x,t) = 0*. The relation between the radius *d* of the channel and *M* is *d =* ℏ*/cM*, for *M ≠ 0, P ≠ 0*, where ℏ= *h*/2*π*; *h* is the Planck constant and *c* the speed of light.

The e.m. field propagation is thus confined within such channels in the polarized medium. Outside them its intensity is zero. The transverse field gradient on the channel boundary acts on the atoms or molecular aggregates in the region surrounding the channels as a selective force given by ^[2,14]^

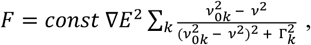

where E is the electric field, *v* its frequency, Γ_*k*_ the damping of the molecular oscillations of frequency *v*_*0k*_. We see that the force is relevant for high field gradient and if the resonant condition *v*_*0k*_ ≃ *v* is met. Moreover, the force *F* is repulsive or attractive depending on the ± sign of the difference 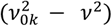, respectively, which also shows that small changes in the frequencies near the resonant values, by producing switching between the ± signs, induce switching between repulsive and attractive forces, respectively. Furthermore, the field frequency within the channel may be perturbed by the presence of the molecule(s) previously attracted on the channel boundary, thus making the force to be attractive (or repulsive) with respect to a molecule with different *v*_*0k*_. The result is that different molecules are selectively collected on the channel boundary. If such molecules may form stable chemical bounds, a dynamical polymerization process is obtained and the polymeric structure coating the channel survives also in the case that the e.m. field vanishes. In the opposite case of low chemical affinity among the collected molecules, the molecular coating will disassemble with the vanishing of the e.m. field. The molecular coating of the field channels may thus represent the observed process of dynamical formation of network branches in the cell cytoskeleton and in cell to cell correlations, affecting their mobility and contractility (cf. the text and Figs. 1,3, column V; S2, Movies S1 and S2). In the case of a fraction α of oriented water molecular dipoles, it can be shown ^[2,3]^ that *M =13,60/*√*α* eV and *d = 146/*√*α* Å, to be compared with *125* Å, the observed average radius of the cytoplasm microtubules. This suggests that *α ≃ 1,4*, i.e. that almost all the dipoles are correlated. Note that *E = 13,60* eV is the hydrogen ionization energy, which may be considered as a threshold energy. A photon of energy *hv > 13,60* eV may propagate with a destructive effect in the coherent dipole ordered region (with possible hydrogen ionization events), thus restoring the spherical symmetry with loss of the filamentary e.m. propagation. On the other hand, a photon of lower energy may be not able to penetrate the ordered patterns. Small amounts of supplied energy may contribute instead to the increase of the polarization, being thus ‘stored’ within the system till a convenient accumulated amount may be used for metabolic activity (e.g. the energy required for a specific chemical reaction to occur), or the threshold *13,60* eV will be reached and then radiated in ionization processes or to open a channel through which propagate (delayed propagation and radiation) ^[2,3,15]^. The possibility of ‘non-thermalizing’ energy storage is thus offered by coherence. In Fig. 1, the panel B-II shows that in the presence of the (positive) energy supply stored in the environment, mitochondrial activity (finalized to supply energy) is resting with respect to the activity shown in panel A-II (absence of sounds; control system).

The mass acquired by the e.m. field produces also a longitudinal component (beside the transvers one) ^[3]^. This component acts as a force directed along the channel pushing the collected molecules, thus, depending on their chemical affinity, either promoting their chemical interaction, or, producing disruption of the microtubule (especially in its ending part, similarly to what observed in the microtubule treadmilling). On the other hand, it is observed that most of the metabolic reactions occurs on the microtubule structures, which also acts as a sort of transport rail system of the cell. We then may conclude that the transverse (attractive or repulsive) force may produce the sequential ordering of interlocked chemical reactions on the microtubule, while its longitudinal component contributes to the chemical transport cell system.

The processes described above crucially depend on the time dependent polarization density *P(x,t)* produced in the cell and in the inter-cellular bath under the action of the external stimuli. Too much high polarization density and its persistence for a too long time interval may prevent the e.m. field disturbance to penetrate through the ordered medium in the described filamentary fashion and the subsequent described processes will not occur. On the contrary, too much low *P(x,t)* and its too short life-time might allow to an even low energy stimulus to fully destroy any polarization pattern. We thus see that the cell functional activity may occur within a window limited by high and low values of the ordering produced by external stimuli.

Thermal effects are among the perturbing agents of the working conditions. Ordered patterns are described in many-body physics in terms of condensation in the ground state of the NG quanta of the long range correlation waves (the dwq in the present case), as mentioned above. In the approximation that the condensed quanta behave as free particles in such a ground state, we have ^[2]^:

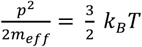

where *T* is the temperature, *p* the momentum, *k*_*B*_ the Boltzmann constant and *m*_*eff*_ the effective mass of the dwq due to the finite linear size *R =* ℏ*/cm*_*eff*_ of the coherent domain. Use of the de Broglie relation *p = h/λ*, in the stationary condition *R = n λ/2*, with *n* an integer, gives

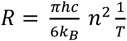

At *T= 300* K, for *n = 1, R = 25* micron, which agrees with the order of magnitude of the observed structures (see the scale reported in the Figures in the text). The above relation shows that in order to keep *T* constant, the system reacts (opposes) to, e.g., an increase of *T*, due for example to an external supply of energy, by increasing the linear size *R* of the coherent correlated region, i.e. shifting toward longer wave lengths correlations (for fixed *n*). Similarly, it opposes to a decrease of *T* by lowering the linear size *R* of the correlated region. We have a dynamical reaction of the system aimed to preserve its functionality against thermal effects.

On the other hand, by considering the system in a given ordered state, i.e. with a given density N of condensed dwq at a stationary temperature ((almost) constant *T*), the minimization of the free energy *F* characterizes the system stationary state: *dF = dU – k*_*B*_*T dS = 0*, with *U* the internal energy and *S* the entropy. We then obtain ^[16,17]^: *dU* ∝ *dN/dt* ∝ *k*_*B*_*TdS*, where as usual we may define heat as *dQ* = *k*_*B*_*T dS*, and which shows that a supply of energy so to have *dU > 0*, implies *dS > 0* and an increase of the rate of change of *dN/dt*, i.e. an *increase in the loss* of condensed dwq, i.e. of correlations in the ground state, since *dS > 0* implies loss of ordering. Thus we see the ‘disordering effect’ of the ‘negative’ sound releasing energy to the system (*dU > 0*) in a ‘non-resonant’ way, possibly leading to the cell explosion. On the contrary, ‘positive’ sounds producing ordering imply *dS < 0*, i.e. *dU < 0* and *dN/dt* < *0*, i.e. a *reduction of the loss* of correlations. We also see the positive effect of dissipating energy by the system (*dU < 0*), provided that a critical threshold is respected.

Another dynamical defense of the system against thermalization is due to the dynamical nonlinearity. The system is made by a large number *N* of components. Their coupling to phonon and to e.m. field gets enhanced by a factor √*N* ^[18]^. In the limit of large *N*, the collective interaction time scale is then shorter by a factor *1/*√*N* than the short range interaction time scale among the individual components. The collective interaction is thus protected against thermal fluctuations. These may interfere with the coherent dynamics only provided that *k*_*B*_*T* is comparable or larger than the collective interaction energy. This guaranties the character of collective dynamics for the system as a whole, a result out of reach in a purely atomistic view and approach.

Finally, we mention that there is an isomorphism between coherent states and self-similarity fractal properties ^[19,20]^ so that we expect that a possible difference between ‘positive’ and ‘negative’ signals may consist in their self-similar fractal properties, either in their different fractal dimension, or in having or not having fractal structure, the ‘positive’ and the ‘negative’ ones, respectively, so that they can induce or not, correspondingly, coherent state configurations with similar fractal properties in the biological matter. A suggestion in such a direction comes to us from the plot in Fig. A3, which represents the signal of the sound used in meditation (one of the “positive” sounds, cf. its effects in Fig. S2). The linear fitting of the log of power (on the ordinate axis) *vs* the log of frequency (on the absciss axis) shows the self-similarity structure of the signal with fractal dimension *d* = *-2,14787E7* (the slope of the fitting straight line). It is in our future plan to analyze the fractal properties, if any, of the other used signals.

In conclusion, the QFT model seems to account for the observed effects that are produced by the diverse sounds on the biological matter. Sounds may perturb the biological functional activity by enhancing or inhibiting in the way described above the coherent dipole dynamics, the formation and propagation of solitons and electrosolitons and the associated nondissipative transport on protein chains of energy and charges trapped by them, the self-focusing propagation of the e.m. field, the formation and depletion of microtubules and of links in the inter-cellular network.

---

**Fig. A1.**
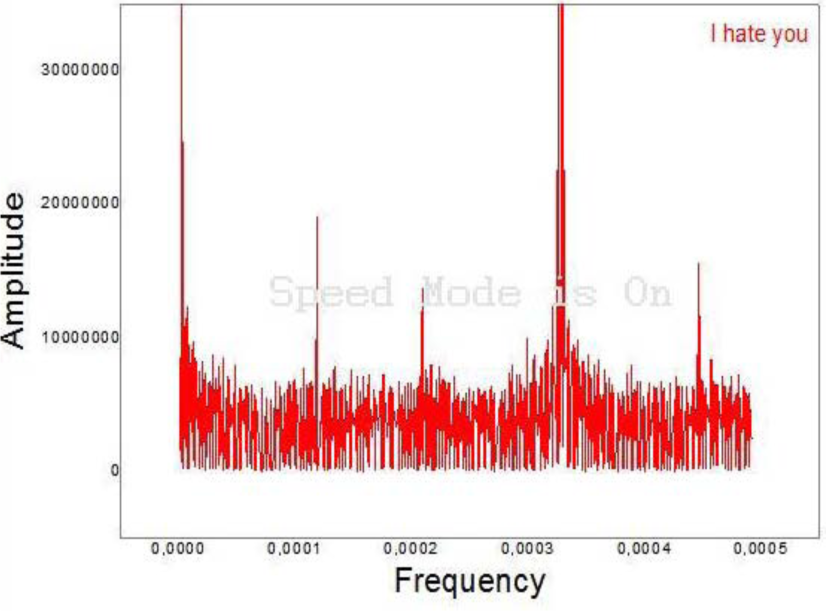
Negative sound (“Ti odio”-I hate you). Signal FFT (finite Fourier transform).

**Fig. A2.**
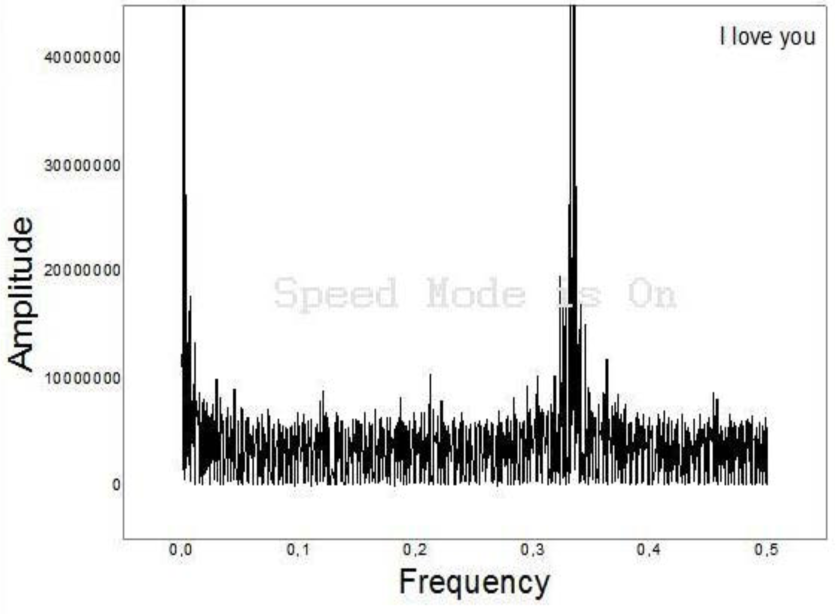
Positive sound (“Ti amo”- I love you). Signal FFT (finite Fourier transform).

**Fig. A3.**
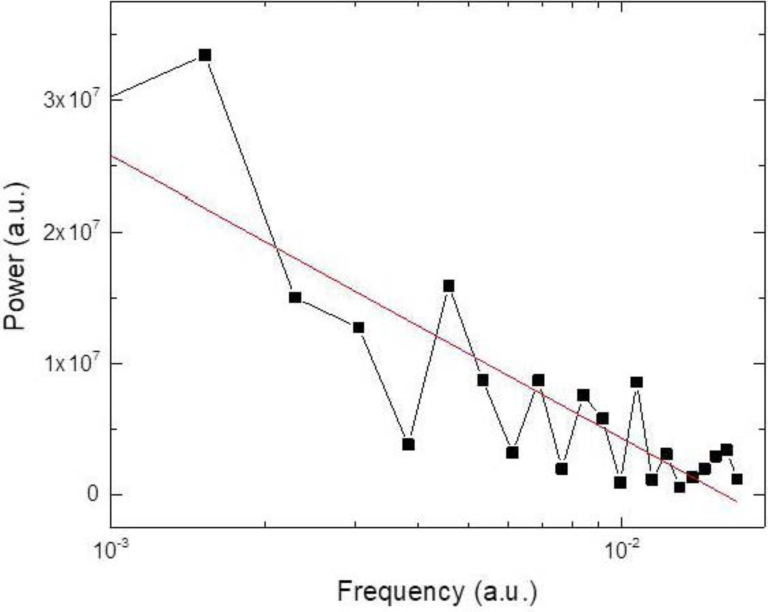
Positive sound (Sound used in meditation). Log-log plot (Power *vs* frequency, arbitrary units). Fractal dimension (straight line slope) d = −2,14787E7 (signal analyzed by use of Origin data analysis software).

## References

1. Albrecht-Buehler G. Rudimentary form of cellular ‘vision’. Proc Natl Acad Sci U S A 1992;89:8288 [PMID: 1518860 DOI: 10.1073/pnas.89.17.8288]

2. Albrecht-Buehler G. A long-range attraction between aggregating 3T3 cells mediated by near-infrared light scattering. Proc Natl Acad Sci U S A 2005;[PMID: 15790680 DOI: 10.1073/pnas.0407763102]

3. Sahu S, Ghosh S, Fujita D, Bandyopadhyay A. Live visualizations of single isolated tubulin protein self-assembly via tunneling current: effect of electromagnetic pumping during spontaneous growth of microtubule. Sci Rep 2014;4:7303 [PMID: 25466883 DOI: 10.1038/srep07303]

4. Sahu S, Ghosh S, Hirata K, Fujita D, Bandyopadhyay A. Multi-level memory-switching properties of a single brain microtubule. Appl Phys Lett 2013;[DOI: 10.1063/1.4793995]

5. Havelka D, Cifra M, Kučera O, Pokorný J, Vrba J. High-frequency electric field and radiation characteristics of cellular microtubule network. J Theor Biol 2011;286:31–40 [PMID: 21782830 DOI: 10.1016/j.jtbi.2011.07.007]

6. Pelling AE, Sehati S, Gralla EB, Valentine JS, Gimzewski JK. Local nanomechanical motion of the cell wall of Saccharomyces cerevisiae. Science 2004;305:1147–50 [PMID: 15326353 DOI: 10.1126/science.1097640]

7. Haase K, Pelling AE. Investigating cell mechanics with atomic force microscopy. J R Soc Interface 2015;12:20140970 [PMID: 25589563 DOI: 10.1098/rsif.2014.0970]

8. Gimzewski JK, Pelling AE, Ventura C. Nanomechanical characterization of cellular activity. 2008;International Publication Number WO 2008/105919 A3

9. Uzer G, Thompson WR, Sen B, Xie Z, Yen SS, Miller S, Bas G, Styner M, Rubin CT, Judex S, Burridge K, Rubin J. Cell Mechanosensitivity to Extremely Low-Magnitude Signals Is Enabled by a LINCed Nucleus. Stem Cells 2015;33:2063–76 [DOI: 10.1002/stem.2004]

10. Rubin C, Turner AS, Bain S, Mallinckrodt C, McLeod K. Low mechanical signals strengthen long bones. Nature 2001, 412:603–4 [PMID: 11493908 DOI: 10.1038/35088122]

11. Rubin CT, Capilla E, Luu YK, Busa B, Crawford H, Nolan DJ, Mittal V, Rosen CJ, Pessin JE, Judex S. Adipogenesis is inhibited by brief, daily exposure to high-frequency, extremely low-magnitude mechanical signals. Proc Natl Acad Sci U S A 2007;104:17879–84 [PMID: 17959771 DOI: 10.1073/pnas.0708467104]

12. Ozcivici E, Luu YK, Adler B, Qin Y-X, Rubin J, Judex S, Rubin CT. Mechanical signals as anabolic agents in bone. Nat Rev Rheumatol 2010;6:50–9 [PMID: 20046206 DOI: 10.1038/nrrheum.2009.239]

13. Facchin F, Bianconi E, Canaider S, Basoli V, Biava PM, Ventura C. Tissue Regeneration without Stem Cell Transplantation: Self-Healing Potential from Ancestral Chemistry and Physical Energies. Stem Cell Int 2018;ID 7412035:8 [DOI: 10.1155/2018/7412035]

14. Martens EA, Thutupalli S, Fourrière A, Hallatschek O. Chimera states in mechanical oscillator networks. Proc Natl Acad Sci U S A 2013;110:10563–7 [PMID: 23759743 DOI: 10.1073/pnas.1302880110]

15. Matsuhashi M, Pankrushina AN, Takeuchi S, Ohshima H, Miyoi H, Endoh K, Murayama K, Watanabe H, Endo S, Tobi M, Mano Y, Hyodo M, Kobayashi T, Kaneko T, Otani S, Yoshimura S, Harata A, Sawada T. Production of sound waves by bacterial cells and the response of bacterial cells to sound. J Gen Appl Microbiol 2005;44:49–55 [DOI: 10.2323/jgam.44.49]

16. Babayi T, Riazi GH. The Effects of 528 Hz Sound Wave to Reduce Cell Death in Human Astrocyte Primary Cell Culture Treated with Ethanol. J Addict Res Ther 2017;08 [DOI: 10.4172/2155-6105.1000335]

17. Lestard NR, Capella MAM. Exposure to Music Alters Cell Viability and Cell Motility of Human Nonauditory Cells in Culture. Evidence-based Complement Altern Med 2016;[PMID: 27478480 DOI: 10.1155/2016/6849473]

18. Lestard NDR, Valente RC, Lopes AG, Capella MAM. Direct effects of music in non-auditory cells in culture. Noise Health 2013;15:307–14 [PMID: 23955127 DOI: 10.4103/1463-1741.116568]

19. Lenzi P, Frenzili G, Gesi M, Ferrucci M, Lazzeri G, Fornai F, Nigro M. DNA damage associated with ultrastructural alterations in rat myocardium after loud noise exposure. Environ Health Perspect 2003;111:467–71 [DOI: 10.1289/ehp.5847]

20. Antunes E, Borrecho G, Oliveira P, de Matos APA, Brito J, Águas A, Martins dos Santos J. Effects of low-frequency noise on cardiac collagen and cardiomyocyte ultrastructure: An immunohistochemical and electron microscopy study. Int J Clin Exp Pathol 2013;6:2333–41

21. Uchiyama M, Jin X, Zhang Q, Hirai T, Amano A, Bashuda H, Niimi M. Auditory stimulation of opera music induced prolongation of murine cardiac allograft survival and maintained generation of regulatory CD4+CD25+ cells. J. Cardiothorac. Surg. 2012;7:26 [PMID: 22445281 DOI: 10.1186/1749-8090-7-26]

22. Ventura C, Graves M, Bergonzoni A, Tassinari R, Cavallini C. Cell melodies : when sound speaks to stem cells. CellR4 2017;5:e2331

23. Naseer SM, Manbachi A, Samandari M, Walch P, Gao Y, Zhang YS, Davoudi F, Wang W, Abrinia K, Cooper JM, Khademhosseini A, Shin SR. Surface acoustic waves induced micropatterning of cells in gelatin methacryloyl (GelMA) hydrogels. Biofabrication 2017;9:015020 [PMID: 28195834 DOI: 10.1088/1758-5090/aa585e]

24. Guo F, Li P, French JB, Mao Z, Zhao H, Li S, Nama N, Fick JR, Benkovic SJ, Huang TJ. Controlling cell– cell interactions using surface acoustic waves. Proc Natl Acad Sci 2015;112:43–8 [PMID: 25535339 DOI: 10.1073/pnas.1422068112]

25. Havelka D, Kučera O, Deriu MA, Cifra M. Electro-acoustic behavior of the mitotic spindle: a semi-classical coarse-grained model. PLoS One 2014;9:e86501.[PMID: 24497952 DOI: 10.1371/journal.pone.0086501]

26. Kučera O, Havelka D, Cifra M. Vibrations of microtubules: Physics that has not met biology yet. Wave Motion 2017;72:13–22 [DOI: 10.1016/j.wavemoti.2016.12.006]

27. Wang N, Tytell JD, Ingber DE. Mechanotransduction at a distance: Mechanically coupling the extracellular matrix with the nucleus. Nat. Rev. Mol. Cell Biol. 2009;10:75–82 [PMID: 19197334 DOI: 10.1038/nrm2594]

28. Guzun R, Karu-Varikmaa M, Gonzalez-Granillo M, Kuznetsov A V., Michel L, Cottet-Rousselle C, Saaremäe M, Kaambre T, Metsis M, Grimm M, Auffray C, Saks V. Mitochondria-cytoskeleton interaction: Distribution of β-tubulins in cardiomyocytes and HL-1 cells. Biochim Biophys Acta - Bioenerg 2011;1807:458–69 [DOI: 10.1016/j.bbabio.2011.01.010]

29. Kuznetsov A V., Javadov S, Guzun R, Grimm M, Saks V. Cytoskeleton and regulation of mitochondrial function: The role of beta-tubulin II. Front Physiol 2013;4 APR [DOI: 10.3389/fphys.2013.00082]

30. Chang CT, Bostwick JB, Steen PH, Daniel S. Substrate constraint modifies the Rayleigh spectrum of vibrating sessile drops. Phys Rev E - Stat Nonlinear, Soft Matter Phys 2013;88:23015 [DOI: 10.1103/PhysRevE.88.023015]

31. Kučera O, Havelka D. Mechano-electrical vibrations of microtubules-Link to subcellular morphology. BioSystems 2012;109:346–55 [DOI: 10.1016/j.biosystems.2012.04.009]

32. Misawa T, Takahama M, Kozaki T, Lee H, Zou J, Saitoh T, Akira S. Microtubule-driven spatial arrangement of mitochondria promotes activation of the NLRP3 inflammasome. Nat Immunol 2013;14:454–60 [PMID: 23502856 DOI: 10.1038/ni.2550]

33. Peleg B, Disanza A, Scita G, Gov N. Propagating cell-membrane waves driven by curved activators of actin polymerization. PLoS One 2011;6:1–11 [DOI: 10.1371/journal.pone.0018635]

34. Dierkes K, Sumi A, Solon J, Salbreux G. Spontaneous Oscillations of Elastic Contractile Materials with Turnover. Phys Rev Lett [Internet] 2014 [cited 2019 Apr 2];113:148102 [DOI: 10.1103/PhysRevLett.113.148102]

35. Bányai LA. A Compendium of Solid State Theory. Springer International Publishing; 2018.

36. Maldovan M. Sound and heat revolutions in phononics. Nature 2013;503 [DOI: 10.1038/nature12608]

37. Meijer DKF, Geesink HJH. Guided Folding of Life’s Proteins in Integrate Cells with Holographic Memory and GM-Biophysical Steering. Open J Biophys 2018;08:117–54 [DOI: 10.4236/ojbiphy.2018.83010]

38. Meijer DKF, Geesink JH. Phonon Guided Biology. Architecture of Life and Conscious Perception Are Mediated by Toroidal Coupling of Phonon, Photon and Electron Information Fluxes at Discrete Eigenfrequencies. NeuroQuantology 2016;14 [DOI: 10.14704/nq.2016.14.4.985]

39. Acbas G, Niessen KA, Snell EH, Markelz AG. Optical measurements of long-range protein vibrations. Nat Commun 2014;5:3076 [DOI: 10.1038/ncomms4076]

40. Gerlich S, Eibenberger S, Tomandl M, Nimmrichter S, Hornberger K, Fagan PJ, Tüxen J, Mayor M, Arndt M. Quantum interference of large organic molecules. Nat Commun 2011;2:263 [PMID: 21468015 DOI: 10.1038/ncomms1263]

41. Zhang S, Cheng J, Qin YX. Mechanobiological modulation of cytoskeleton and calcium influx in osteoblastic cells by short-term focused acoustic radiation force. PLoS One 2012;7:e38343.[DOI: 10.1371/journal.pone.0038343]

42. Kamkin A, Kiseleva I. Mechanosensitivity and Mechanotransduction. Springer Netherlands; 2011.

43. Mavromatos NE. Non-linear dynamics in biological microtubules: Solitons and dissipation-free energy transfer. In: Journal of Physics: Conference Series. IOP Publishing; 2017.

44. Manka R, Ogrodnik B. A model of soliton transport along microtubules. J Biol Phys 1991;18:185–9 [DOI: 10.1007/BF00417807]

45. Abdalla E, Maroufi B, Melgar BC, Sedra MB. Information transport by sine-Gordon solitons in microtubules. Phys A Stat Mech its Appl 2001;301:169–73 [DOI: 10.1016/S0378-4371(01)00399-5]

46. Liu D, Every AG, Tománek D. Long-wavelength deformations and vibrational modes in empty and liquid-filled microtubules and nanotubes: A theoretical study. Phys Rev B 2017;95:205407 [DOI: 10.1103/PhysRevB.95.205407]

47. Prodan E, Prodan C. Topological Phonon Modes and Their Role in Dynamic Instability of Microtubules. Phys Rev Lett 2009;103:248101 [DOI: 10.1103/PhysRevLett.103.248101]

48. Prodan E, Dobiszewski K, Kanwal A, Palmieri J, Prodan C. Dynamical Majorana edge modes in a broad class of topological mechanical systems. Nat Commun 2017;8:14587 [DOI: 10.1038/ncomms14587]

49. Aghaei A, Dayal K, Elliott RS. Symmetry-adapted phonon analysis of nanotubes. J Mech Phys Solids 2013;61:557–78 [DOI: 10.1016/j.jmps.2012.09.008]

50. Zandonella C. Dying cells dragged screaming under the microscope. Nature 2003;423:106–7 [DOI: 10.1038/423106b]

51. Dal Lin C, Marinova M, Rubino G, Gola E, Brocca A, Pantano G, Brugnolo L, Sarais C, Cucchini U, Volpe B, Cavalli C, Bellio M, Fiorello E, Scali S, Plebani M, Iliceto S, Tona F. Thoughts modulate the expression of inflammatory genes and may improve the coronary blood flow in patients after a myocardial infarction. J Tradit Complement Med 2018;8:150–63 [PMID: 29322004 DOI: 10.1016/j.jtcme.2017.04.011]

## References for appendix 1

1. Del Giudice E, Doglia S, Milani M, Vitiello G. A quantum field theoretical approach to the collective behavior of biological systems. Nucl Phys B 1985; 251(FS 13): 375–400.

2. Del Giudice E, Doglia S, Milani M, Vitiello G. Electromagnetic field and spontaneous symmetry breakdown in biological matter. Nucl. Phys. B 1986; 275(FS 17):185–199.

3. Del Giudice E, Doglia S, Milani M, Vitiello G. Structure, correlations and electromagnetic interactions in living matter: Theory and applications. In Biological coherence and response to external stimuli, H. Fröhlich Ed., Springer-Verlag, Berlin, 1988; 49–64.

4. Goldstone J, Salam, A, Weinberg, S. Broken Symmetries, Phys. Rev 1962; 127: 965–970.

5. Umezawa H. Advanced field theory: Micro, macro, and thermal physics. New York, AIP, 1993.

6. Blasone M, Jizba P, Vitiello G. Quantum Field Theory and its macroscopic Manifestations, Imperial College Press, London, 2011.

7. Davydov A S. Solitons in Molecular System. Dordrecht, Reidel, 1985.

8. Brizhik L, Cruzeiro-Hansson L, Eremko A, Olkhovska Yu. Soliton dynamics and Peierls-Nabarro barrier in a discrete molecular chain. Phys Rev B 2000; 61(2): 1129–1141.

9. Brizhik L. Influence of electromagnetic field on soliton mediated charge transport in biological systems. Electromagn Biol Med 2015;34(2): 123–132 (ID: 1036071; DOI:10.3109/15368378.2015.1036071).

10. McDermott M L, Vanselous H, Corcelli S A, PetersenP B. DNA’s Chiral Spine of Hydration. ACS Cent Sci 2017; 3 (7) : 708–714.

11. Marburgher J H. Self-focusing. Theory Prog Quant Electr 1975; 4:35–110.

12. Chiao R Y, Gustafson T K and Kelley P L. Self-Focusing of Optical Beams. In: Boyd, R. W., Lukishova, S G. and Shen, Y R. (Eds.), Self Focusing: Past and Present. Springer, New York, 2009; 129–143.

13. Zakharov V E, Shabat A B. Exact theory of two-dimensional self-focusing and automodulation of waves in nonlinear media. Sov J Exp Theor Phys 1971; 61:118–134.

14. Askar’yan G A. The self-focusing effect. Sov phys Usp 1974;16: 680.

15. Mandoli D F, Briggs W. Optical properties of etiolated plant tissues. Proc Nat Acad Sc USA 1982; 79: 2902–2906.

16. Celeghini E, Rasetti M, Vitiello G. Quantum dissipation. Annals of Physics (N.Y.) 1992; 215:156–170.

17. Vitiello G. Dissipation and memory capacity in the quantum brain model. Int J Mod Phys 1995; B9: 973–989.

18. Del Giudice E, Vitiello G. The role of the electromagnetic field in the formation of domains in the process of symmetry breaking phase transitions. Phys Rev A 2006; 74: 02210, 5.

19. Vitiello G. Fractals, coherent states and self-similarity induced noncommutative geometry. Phys. Lett. A 2012; 376: 2527–2532.

20. Vitiello G. Coherent states, fractals and brain waves. New Mathematics and Natural Computing 2009; 5: 245–264.

